# ProtPipe: A Multifunctional Data Analysis Pipeline for Proteomics and Peptidomics

**DOI:** 10.1101/2023.12.12.571327

**Authors:** Ziyi Li, Cory A. Weller, Syed Shah, Nicholas Johnson, Ying Hao, Jessica Roberts, Colleen Bereda, Sydney Klaisner, Pedro Machado, Pietro Fratta, Leonard Petrucelli, Mercedes Prudencio, Björn Oskarsson, Nathan P. Staff, Dennis W. Dickson, Mark R. Cookson, Michael E. Ward, Andrew B. Singleton, Mike A. Nalls, Yue A. Qi

## Abstract

Mass spectrometry (MS) is a technique widely employed for the identification and characterization of proteins, personalized medicine, systems biology and biomedical applications. By combining MS with different proteomics approaches such as immunopurification MS, immunopeptidomics, and total protein proteomics, researchers can gain insights into protein-protein interactions, immune responses, cellular processes, and disease mechanisms. The application of MS-based proteomics in these areas continues to advance our understanding of protein function, cellular signaling, and complex biological systems. Data analysis for mass spectrometry is a critical process that includes identifying and quantifying proteins and peptides and exploring biological functions for these proteins in downstream analysis. To address the complexities associated with MS data analysis, we developed ProtPipe to streamline and automate the processing and analysis of high-throughput proteomics and peptidomics datasets. The pipeline facilitates data quality control, sample filtering, and normalization, ensuring robust and reliable downstream analysis. ProtPipe provides downstream analysis including identifying differential abundance proteins and peptides, pathway enrichment analysis, protein-protein interaction analysis, and MHC1-peptide binding affinity. ProtPipe generates annotated tables and diagnostic visualizations from statistical postprocessing and computation of fold-changes across pairwise conditions, predefined in an experimental design. ProtPipe is well-documented open-source software and is available at https://github.com/NIH-CARD/ProtPipe, accompanied by a web interface.

## Introduction

Recent advancements in mass spectrometry (MS) have revolutionized the field of high-throughput proteomics, enabling accurate and rapid analysis of thousands of proteins across multiple biological samples[1–5]. This enhanced capability has allowed researchers to characterize the proteome on an unprecedented scale and to investigate dynamic changes in protein expression under various experimental conditions[1, 6–8]. Although multiplex barcoding MS (e.g., tandem mass tag) enhances throughput, it also results in more complex samples, often requiring offline fractionation and resulting in batch effects. Label-free MS has become an indispensable tool in protein quantification, offering advantages in sample preparation, dynamic range, and absolute quantification at the protein level[9]. Its applications span a wide range of research areas, including proteomics, personalized medicine, and biomarker discovery[9, 10]. With continued advancements in MS technology and data analysis strategies, label-free approaches hold great promise for further advancements in understanding complex biological systems.

Protein-protein interactions (PPIs) play crucial roles in cellular processes and disease mechanisms. The integration of affinity purification with mass spectrometry (AP-MS) has revolutionized the study of PPIs in high-throughput proteomics[11]. Following affinity purification, which can be co-immunoprecipitation or proximity-labeling, the isolated protein complexes are subjected to mass spectrometry analysis, enabling the identification and quantification of the interacting partners. Mass spectrometry provides information about protein identities, post-translational modifications, and dynamic changes in the interactome. By selectively isolating and identifying interacting partners, AP-MS allows for the systematic exploration of complex biological networks and dynamic changes in the interactome. This powerful approach has broad applications in understanding cellular processes, disease mechanisms, and drug discovery, contributing to advancements in proteomics research[12–14].

Immunopeptidomics has emerged as a powerful approach to enhance our understanding of antigen presentation, immune recognition, and vaccine development[15, 16]. This technique enables the comprehensive characterization and quantification of peptides presented by major histocompatibility complexes (MHC) on the cell surface. Recent applications of quantitative immunopeptidomes have yielded significant insights into the identification of tumor-specific antigens, characterization of viral epitopes, and exploration of the impact of diseases on the immunopeptidome. These advancements pave the way for personalized immunotherapies, precision medicine, and the development of targeted vaccines against various diseases.

Advancements in proteomics, akin to genomics, have led to an exponential increase in data size and complexity, shifting the bottleneck from the experimental stage to data analysis. Proteomics data post-analysis is a vital phase in transforming raw experimental data into meaningful biological insights. A multitude of software tools and algorithms are available to aid researchers in processing, analyzing, and interpreting complex proteomics datasets. However, many of these tools are designed for genetics and genomics studies, leading to statistical bias. Surprisingly, a handful of integrated bioinformatic tools are not available for MS-based proteomics. This article presents ProtPipe, a comprehensive data analysis tool, outlining its key features and methodology. We first describe a comprehensive exposition of ProtPipe’s essential features accompanied by a detailed description of the employed data analysis methods. The intricacies of the research methodology are meticulously outlined, offering a lucid understanding of the procedures involved. Furthermore, an extensive evaluation of the performance and applicability is conducted by subjecting it to the analysis of three proteome-wide quantification data sets.

These data sets, meticulously generated with diverse experimental designs, serve as the basis for evaluating ProtPipe’s robustness and adaptability across varying scenarios. By subjecting ProtPipe to this rigorous assessment, we ascertain its ability to effectively handle the complexities inherent in diverse experimental conditions. The ensuing discussion delves into the obtained results, with an emphasis on comparative analysis and biological relevance. The findings contribute to establishing ProtPipe as a valuable and versatile tool for proteomics research, poised to advance our understanding of intricate biological phenomena that are easily deployable in many environments.

This repository serves as a valuable resource for researchers seeking to engage in MS-based proteomics including the most common data acquisition modes, such as data independent acquisition (DIA) and data dependent acquisition (DDA). Our workflow provides an easy, efficient, and reproducible, even with minimal command line experience. A user-friendly wrapper script executes DIA-NN[17] within a pre-built Singularity image, allowing users to swiftly export protein abundance from raw MS output. Notably, the script accommodates protein abundance estimates from DIA-NN and Spectronaut, and FragPipe[18–20], enhancing the versatility of the workflow. Moreover, the repository includes a preconfigured R environment, which enables seamless post-processing of the protein abundance estimates. Researchers can utilize this environment to generate comprehensive quality control reports, perform diverse analyses, and create visualizations, all within an accessible framework (Figure 1).

**Figure 1.**
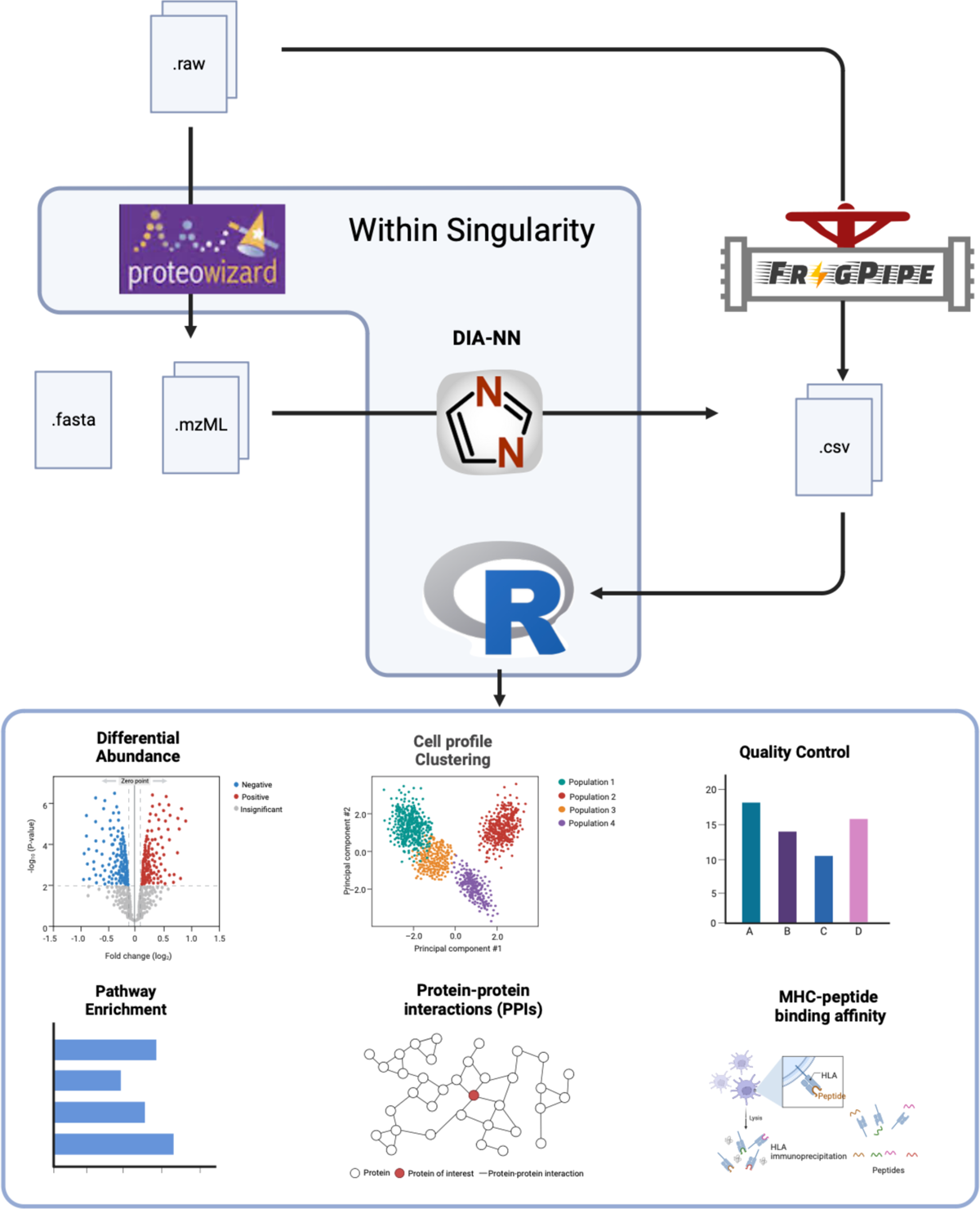
The comprehensive workflow of ProtPipe, a multifunctional data analysis pipeline for proteomics and peptidomics. The database search is completed by DIA-NN in our pre-built singularity or offline Spectronauts for DIA, and offline Fragpipe for DDA. The output csv files (i.e., protein group or peptides) are analyzed by Protpipe which generates figures and datasets for quality control, clustering analysis, differential abundance analysis, pathway enrichment analysis, protein-protein interaction networks, and immunopeptides binding affinity.

## Materials and Methods

### ProtPipe Feature Overview

1. **Ease of use and reproducibility**. Scientific advancement suffers when experimental results are not reproducible and when computational analysis is complicated to perform or locked behind proprietary software. ProtPipe facilitates reproducibility by operating within a Singularity container where all dependencies are pre-installed. As a result, analyses will produce identical results for a given set of inputs no matter the host operating system configuration. Because all required tools come pre-installed in the container, ProtPipe requires only minimal command-line experience, significantly lowering the barrier to entry for proteomics data analysis. This fully open-source project is hosted at our GitHub repository <u>NIH-CARD/ProtPipe</u>.
2. **Quality control and data preprocessing.** ProtPipe includes flexible quality control steps (with reasonable default settings) for evaluating data reliability and boosting the signal-to-noise ratio prior to conducting analyses. Quality control includes steps such as excluding proteins with globally low abundance and samples with low protein group counts (signifying low sample quality). After removal of problematic proteins or samples, protein abundance must be scaled and normalized to account for systematic variations and technical biases from sample preparation, instrument performance or data acquisition. ProtPipe employs median normalization to adjust protein abundance values across samples. Briefly, we calculate a global median across all unadjusted samples, then apply a sample-specific correction such that the median of each corrected sample matches the original global median. By default, median normalization is achieved by adding a constant to all abundances per sample (we include the option to multiply by a constant if desired).
3. **Peptide and Protein Quantification**. ProtPipe provides researchers with the accurate quantification of peptides across large-scale proteomic datasets DIA-NN. This efficient setup enables researchers to quickly estimate protein abundance from DIA raw output. Additionally, our script accommodates protein abundance estimates obtained from Spectronaut, and FragPipe, thus significantly enhancing the workflow’s versatility. By offering a unified platform to integrate outputs from different software tools, researchers can easily analyze and compare protein abundance, fostering a comprehensive understanding of proteomics data. This distinctive feature significantly augments the workflow’s versatility and fosters a more comprehensive and integrated analysis of proteomics data.
4. **Differential Intensity Analysis.** Much like differential gene expression analysis from RNA sequencing experiments, differential intensity analysis identifies proteins for which intensity is significantly associated with an experimental group or treatment. Only proteins detected more than half of all biological replicates could be used for differential intensity analysis. We test for significant differential protein expression using the t-test on the scaled and normalized data, followed by fold change analysis and multiple comparisons correction by the Benjamini-Hochberg procedure. For labeling differential intensity gene candidates, ProtPipe defaults to a log2 fold-change minimum threshold of 2, and p-value threshold of ≤ 0.05 (these thresholds are easily modified). Annotation and enrichment analysis is performed with the R package clusterProfiler[21]. This analysis leverages the Gene Ontology (GO) database [22](http://geneontology.org/) to annotate proteins according to their Biological Process, MolecularFunction, and Cellular Component. Pathway analysis is performed using the Kyoto Encyclopedia of Genes and Genomes (KEGG) database[23]. Enrichment analyses are ranked by protein intensity, with the top enriched entries (default N=20) represented graphically.
5. **Identification of protein-protein Interactions.** ProtPipe analyzes affinity purification with mass spectrometry (AP-MS) data for identification of PPIs, aiding in the understanding of protein function, dysfunction and regulation within biological systems. PPIs are identified using the extensive STRING database (https://string-db.org, [24]) which offers an expansive repository encompassing established and predicted PPIs sourced from a diverse spectrum of experimental data, computational projections, and text mining.
6. **Immunopeptidome deconvolution.** Neoantigen prediction connects genomics and immunology, facilitating treatment tailored to an individual’s genotype. ProtPipe employs MHCflurry, a machine learning tool for predicting potential neoantigens and their binding affinity with the major histocompatibility complex (MHC).

### Mass Spectrophotometry Methods

The sample preparation for mass spectrometry (MS)-based proteomics was performed on our previously published fully automated pipeline (Reilly et al., 2021). The resulting peptides were measured and normalized using an automated colorimetric assay on a Bravo (Agilent Technologies) robot. The final resulting tryptic peptides were separated on a nano column (75 μm × 500 mm, 2 μm C18 particle) using a 2-h efficient linear gradient (Phase B, 2-35% ACN) on an UltiMate 3000 nano-HPLC system. We used data independent acquisition (DIA) discovery proteomics on a hybrid Orbitrap Eclipse mass spectrometer. Specifically, MS1 resolution was set to 120K, and MS2 resolution was set to 30K. For DIA isolation, the precursor range was set to 400–1000 m/z, and the isolation window was 8 m/z, resulting in 75 windows for each scan cycle (3 s). High collision dissociation was used for fragmentation with 30% collision energy. The AGC target was set to 800% for MS2 scan (improve MS2 spectra quality). The MS2 scan range was defined as 145–1450 m/z, which covers the majority of fragment ions of typical tryptic peptides.

The database search was performed at the protein level by Spectronaut software 16.0. We used a library-free module (i.e., direct DIA), which does not require a preconstructed high-quality project-specific spectral library but only uses a proteome sequence file. Specifically, the MS/MS spectra were searched against a proteome reference database containing 17,125 proteins (reviewed) mapped to the mouse reference genome obtained via the UniProt Consortium. All spectra were allowed ‘‘trypsin and lysC’’ enzyme specificity, with up to two missed cleavages. Allowed fixed modifications included carbamidomethylation of cysteine and the allowed variable modifications for whole proteome datasets were acetylation of protein N-termini, oxidized methionine. The false discovery rates of peptide and protein were all set as 1%. Three replicate MS analyses were done from each sample. Further analysis was performed on the mean values of the replicate MS analyses. DESeq2 was implemented to analyze the differential expression between sample types. R software was used to analyze and visualize data.

## Results

### Performance evaluation of ProtPipe

In this section, we present the results of our performance evaluation of ProtPipe, focusing on key metrics essential for efficient data analysis. To provide a comprehensive understanding of ProtPipe’s performance, we meticulously collected and analyzed data on CPU usage, memory consumption, network activity, and disk utilization across various sample sizes and the computational resources allocated for our analytical pipeline (Table 1). Despite the expectation of the database searching stage being primarily limited by CPU resources, our analysis revealed significant CPU-idle time during this stage. The insights gained from this evaluation are instrumental in optimizing ProtPipe for robust and efficient proteomic data analysis.

**Table 1.**
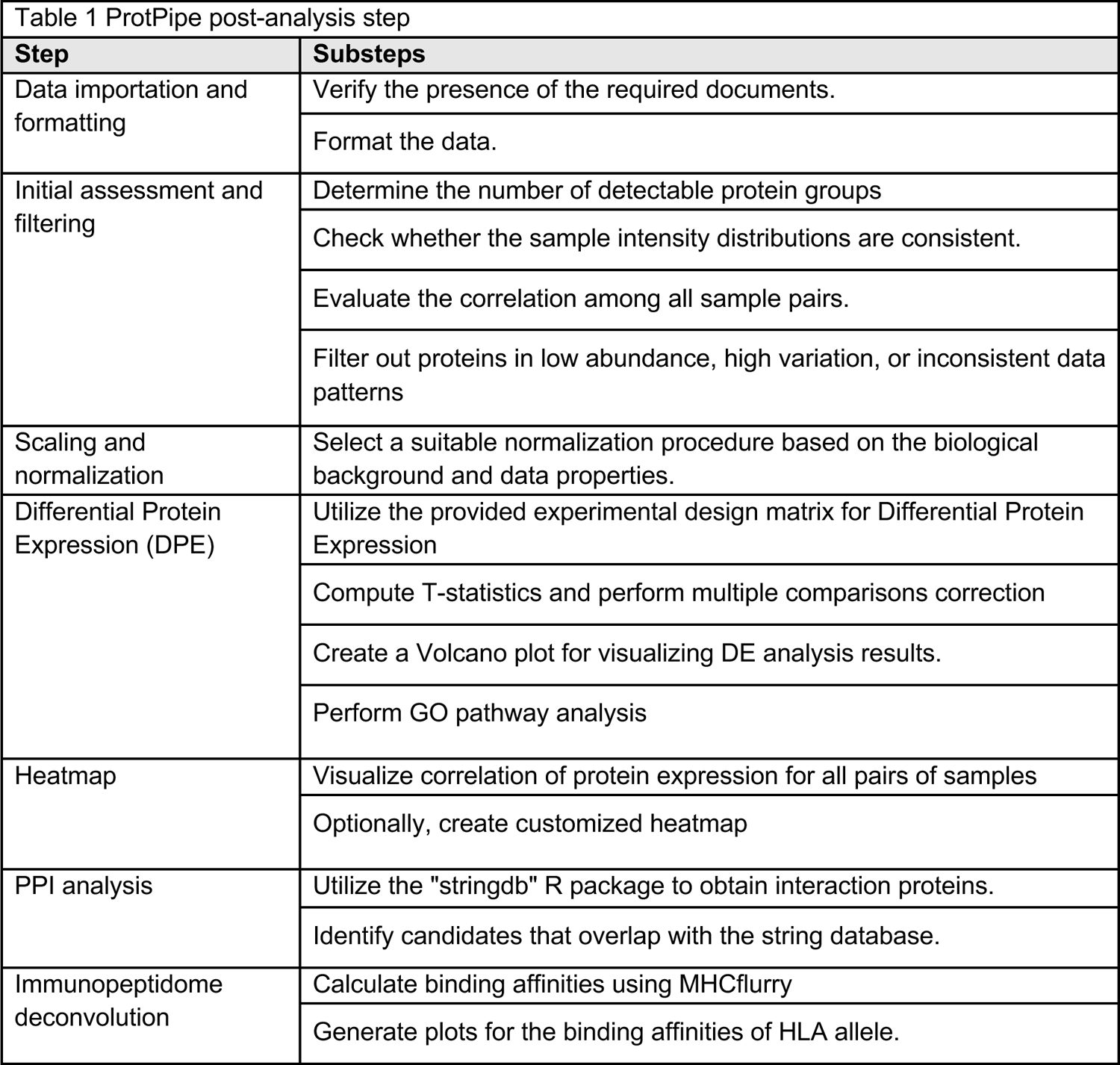
Overview of ProtPipe steps.

**Table 2.**
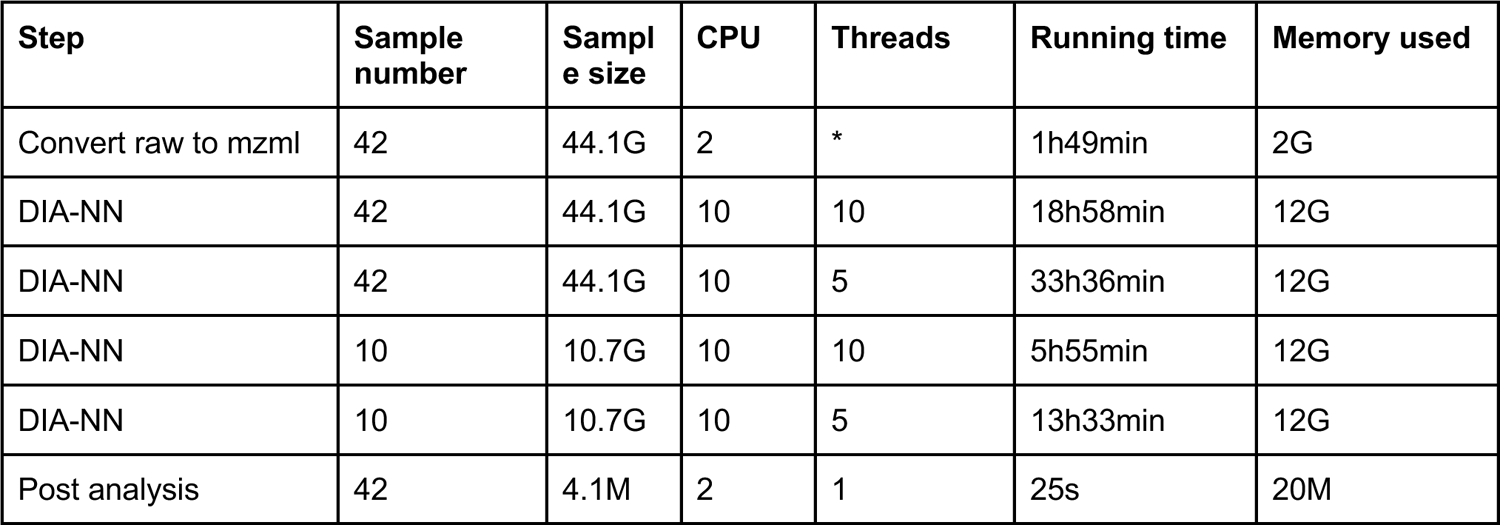
Resource requirements for various stages of data processing with ProtPipe.

| Step | Sample number | Sample size | CPU | Threads | Running time | Memory used |
| --- | --- | --- | --- | --- | --- | --- |
| Convert raw to mzml | 42 | 44.1G | 2 | * | 1h49min | 2G |
| DIA-NN | 42 | 44.1G | 10 | 10 | 18h58min | 12G |
| DIA-NN | 42 | 44.1G | 10 | 5 | 33h36min | 12G |
| DIA-NN | 10 | 10.7G | 10 | 10 | 5h55min | 12G |
| DIA-NN | 10 | 10.7G | 10 | 5 | 13h33min | 12G |
| Post analysis | 42 | 4.1M | 2 | 1 | 25s | 20M |

### Case Study 1: Analysis of a Large-Scale Proteomics Data set

The presented dataset has been generated through the application of the proteomics methodology in characterizing human induced pluripotent stem cells (iPSC) and derived neurons[19]. This method has been conceived as an integral facet of the broader iPSC Neurodegenerative Disease Initiative (iNDI), a concerted and expansive endeavor directed towards the comprehensive elucidation of the conceivable manifestations inherent to hereditary maladies within neural cellular substrates[25].

The proposed analytical pipeline is designed to encompass a comprehensive suite of data quality control measures, including Number of Protein Groups, Distribution of Protein Intensity, and Correlation Among Replicates. Notably, the analysis reveals the detection of a noteworthy count exceeding 7500 proteins across the diverse samples (Figure 1A). This substantial assembly of identified proteins attests to the efficacy of experimental protocols, which have effectively encompassed a wide spectrum of proteins inherent to the samples to the caliber of the data and its propensity for reproducibility. The evaluation of protein intensity values’ distribution (Figure 1B) is conducted to ascertain its uniformity and coherence. A protein intensity profile that is evenly distributed across the samples serves as a compelling indicator of data quality, with its consistency underscored. The manifestation of such uniform distribution signifies the absence of skewness or predisposed biases within the data—an encouraging revelation. In light of this observation, the necessity for data normalization might be rendered redundant, given the apparent absence of conspicuous systematic aberrations necessitating correction. Correlation Analysis, an indispensable analytical technique for discerning interrelationships among variables within a dataset, yields valuable insights. Within this dataset, a discernibly robust correlation exhibited within the replicates serves as a robust testament (Figure 1C).

Cluster analysis stands as a potent analytical method wielded for the discernment of intricate patterns and discernible groupings inherent to intricate datasets. The visualization of these clusters not only serves as an illuminating pictorial representation but also yields profound insights into the fundamental biological dynamics that underlie the data. Within the framework of cluster analysis, our meticulously designed pipeline presents a selection of three distinct cluster methods: Principal Component Analysis (PCA) (Figure 1D), Uniform Manifold Approximation and Projection (UMAP)(Figure S1A), and Hierarchical Clustering (HC)(Figure S1B).

Differential abundance analysis in proteomics involves comparing the abundance of proteins in different experimental conditions. This analysis aims to identify proteins that significantly differ in abundance, potentially providing insights into biological processes, disease mechanisms, or treatment effects. We use the student’s t-test to test for differences in protein abundance while accounting for inherent data variability in proteomics. Based on the provided design matrix, ProtPipe will generate specific comparisons as intended. We illustrate this with an example of the comparison between fully differentiated neurons (on day 28) and iPSCs. During our analysis, the volcano plot serves as a dynamic tool that allows for flexibility in parameter settings, ensuring that the visualization aligns with the nuances of our data. By default, significant differences in abundance require a log2 fold-change of 2 and a False Discovery Rate (FDR) threshold of 0.01 (Figure 1E). Orange and blue color points signify significantly increased or decreased expression in iPSC-derived neurons compared to iPSCs, respectively. These specified threshold values can be adjusted based on the specific context of the investigation.

Furthermore, we understand the importance of highlighting key genes that exhibit significant changes. The top 5 genes that show the most pronounced up- or down-regulation within the comparison would be labeled by default. Specific user-defined genes of interest can also be labeled.

Subsequent to the identification of these differentially abundant proteins, the subsequent endeavor involves elucidating the functional context and implications of these alterations. By delving into the pathways intricately linked to these proteins, a more profound comprehension emerges regarding how these changes may intricately impact cellular functions. Pathway analysis encompasses a comprehensive examination of Gene Ontology (GO) terms, which include Cellular Component (CC) (Figure 2F), Biological Process (BP)(Figure S1C), and Molecular Function (MF)(Figure S1D). These three categories provide a holistic understanding of the functional context of genes and proteins, shedding light on their cellular localization, roles in biological processes, and molecular activities.

**Figure 2.**
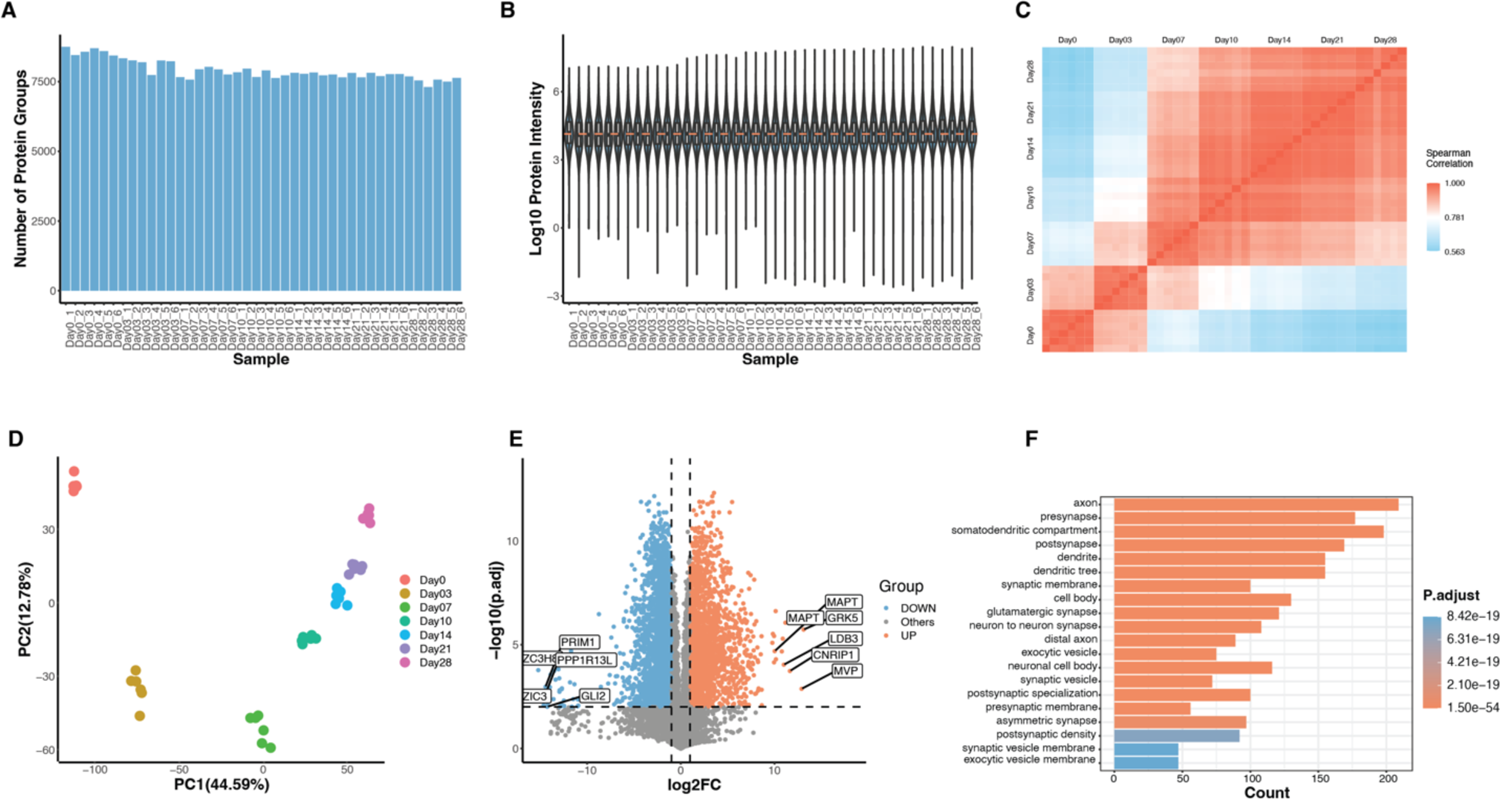
Analysis of a large-scale proteomics data set. A. Distribution of identified protein groups. B. Distribution of protein intensity. C. Correlation among biological replicates. D. Principal Component Analysis (PCA) plot. E. Volcano plot showing differential abundance analysis results between two groups. Orange dots represent upregulated proteins, blue dots indicate downregulated proteins, and gray dots signify proteins with no significant alterations. F. Set of bar charts present Gene Ontology (GO) terms related to Cellular Component (CC).

Progressing in the analysis, the intricate patterns of protein intensity across the samples through the utilization of a heatmap. This visual representation serves as a valuable tool for gaining deeper insights into the dynamics of protein expression, potentially revealing underlying trends and associations. In the default configuration, the heatmap includes all the proteins that have been detected (Figure 3A). This comprehensive view provides a broad perspective of protein expression patterns across the dataset.

**Figure 3:**
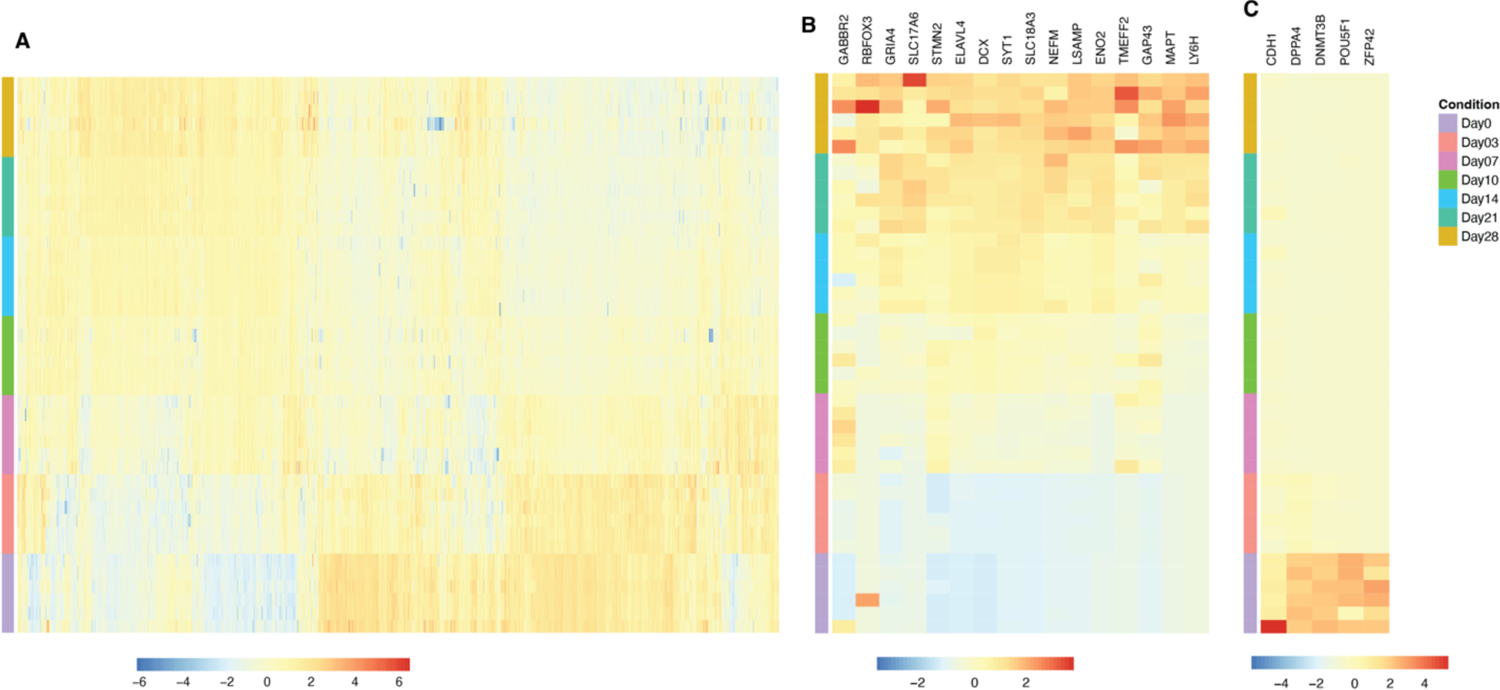
Heatmap analysis of protein intensity Patterns. A. Heatmap patterns of protein intensity across all detected proteins in the samples. B. Customized heatmap generated using a select gene list, focusing on marker genes closely associated with neuron cells. C. Customized heatmap generated using a provided gene list, highlighting marker genes unique to induced pluripotent stem cells (iPSCs).

Additionally, our pipeline offers the flexibility to generate customized heatmaps using gene lists provided by researchers (Figure 3B and 3C). We provide two specific subsets of heatmaps as examples. One set of heatmaps encompasses marker genes closely associated with neuron cells, focusing on marker genes closely associated with neuron cells. This allows researchers to delve into specific protein expression dynamics relevant to neurons. In contrast, the other set of heatmaps are tailored to iPSC marker genes. Researchers can utilize this set to explore gene expression patterns specific to iPSCs.

This versatility in heatmap analysis empowers researchers to conduct more targeted investigations aligned with their research objectives. Whether studying neuron-related gene expression or iPSC-specific dynamics, our pipeline accommodates a wide range of analytical needs.

### Case Study 2: Protein-Protein Interactome Study

Expanding on its proficiency in proteomics data analysis, ProtPipe also possesses the capability to handle AP-MS (Affinity Purification-Mass Spectrometry) data. The dataset featured in this context originates from our previous publication focused on UNC13A, a genetic risk factor associated with amyotrophic lateral sclerosis (ALS) and frontotemporal dementia (FTD)[18]. To identify proteins binding to UNC13A, we conducted a pull-down experiment. During the quality control phase, the number of detected proteins and the distribution of protein abundance within both the negative control group and the UNC13A pull-down group are assessed (Figure 4A, 3B). Notably, the control group exhibited a limited number of detected proteins and lower protein intensity levels. The calculation of sample correlations was undertaken to demonstrate the reproducibility of replicate experiments (Figure 4C).

**Figure 4:**
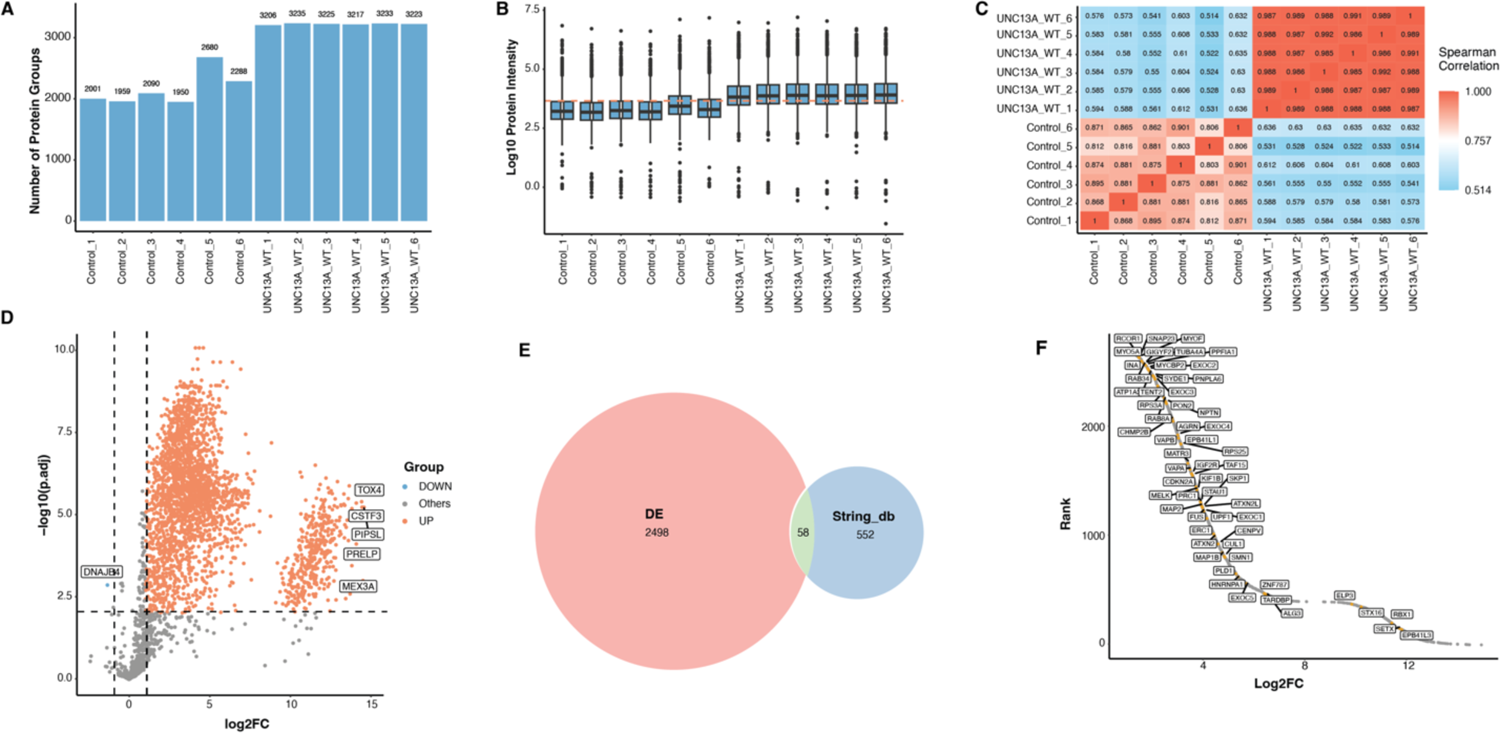
Comprehensive analysis of UNC13A RNA-Binding proteins (RBPs) A-B. Bar chart and box plot depicting the number of detected proteins (A) and the distribution of protein abundance levels (B) in both the control group and the UNC13A pull-down group. C. Reproducibility assessment by replication correlation. D. Identification of Enriched Proteins. Volcano plot illustrating proteins significantly enriched in the presence of UNC13A RNA compared to a negative control. E. Venn diagram showing interacting proteins obtained by leveraging the STRING database and potential interacting proteins compared with experimental data. F. Rank plot of the proteins based on their fold change in abundance. Yellow dots on the plot represent proteins that are previously known to interact with the target protein, as obtained from the STRING database.

We identified several proteins which exhibited a statistically significant enrichment in the presence of UNC13A RNA when contrasted with a negative control RNA (Figure 4D). Furthermore, we conducted GO enrichment analysis focusing on UNC13A RNA-binding proteins (RBPs), revealing that the most prominently enriched biological processes encompassed RNA metabolism, mRNA processing, and RNA splicing (Figure S2A). Additionally, our molecular function analysis demonstrated significant enrichment in terms pertaining to RNA binding (Figure S2B). In terms of cellular compartment analysis, our findings exhibited notable enrichments within the nuclear body and spliceosome categories (Figure S2C).

In investigating PPIs, we leveraged the robust capabilities offered by the STRING database, which serves as a resource for the comprehensive exploration of these intricate molecular associations. In comparison with the data from the STRING database, we observed a noteworthy convergence of potential proteins that may interact with the target protein (Figure 4E). This convergence serves as evidence to substantiate the robustness and reliability of experimental data. Our pipeline serves as a valuable tool for researchers, facilitating the discernment of proteins that have been previously demonstrated to engage in interactions with the target protein in the context of AP-MS experiments (Figure 4F). Moreover, it provides a means by which researchers can explore and unveil novel potential interacting proteins, thereby advancing the depth of understanding in this domain.

### Case Study 3: Identification of allele specific peptides with immunopeptidomics

Our pipeline is designed to assist in the identification of peptides that exhibit higher binding affinities to MHC molecules.In this context, we deployed the pipeline in our previous publication within the context of MS-based proteogenomic profiling, aiming to ascertain potential immunogenic peptides presented by each of human leukocyte antigen (HLA) class I alleles in melanoma and EGFR-mutant lung adenocarcinoma.

The total count of peptides identified within class I immunopeptidome from cancer cell lines and tumors range from 1520 to 3509 (Figure 5A). With HLA genotyping, we deconvoluted the total HLA-peptides to each corresponding HLA allele in PC9 cells (Figure 5B). It is noteworthy that HLA-C07 exhibits the highest peptide count. Subsequently, we delve into an examination of peptides manifesting strong binding affinities with these specific HLA alleles. There are a multitude of peptides that demonstrate the most robust binding affinity with HLA-A02 (Figure 5C). To emphasis on the peptides exhibiting the most potent binding affinities, a ranking plot of peptide-HLA binding affinities with each peptide specific binding to a particular HLA is denoted by distinct colors (Figure 5D). Notably, the top 5 peptides demonstrating the strongest binding affinities are all associated with HLA-A-02. The plot of peptide binding patterns across various HLA alleles highlights the top 5 peptides with the most robust binding affinities for each respective allele (Figure 5E). Furthermore, all MHC peptide binding affinity analyses were conducted in the H1975 cells, as demonstrated in Figure S3.

**Figure 5.**
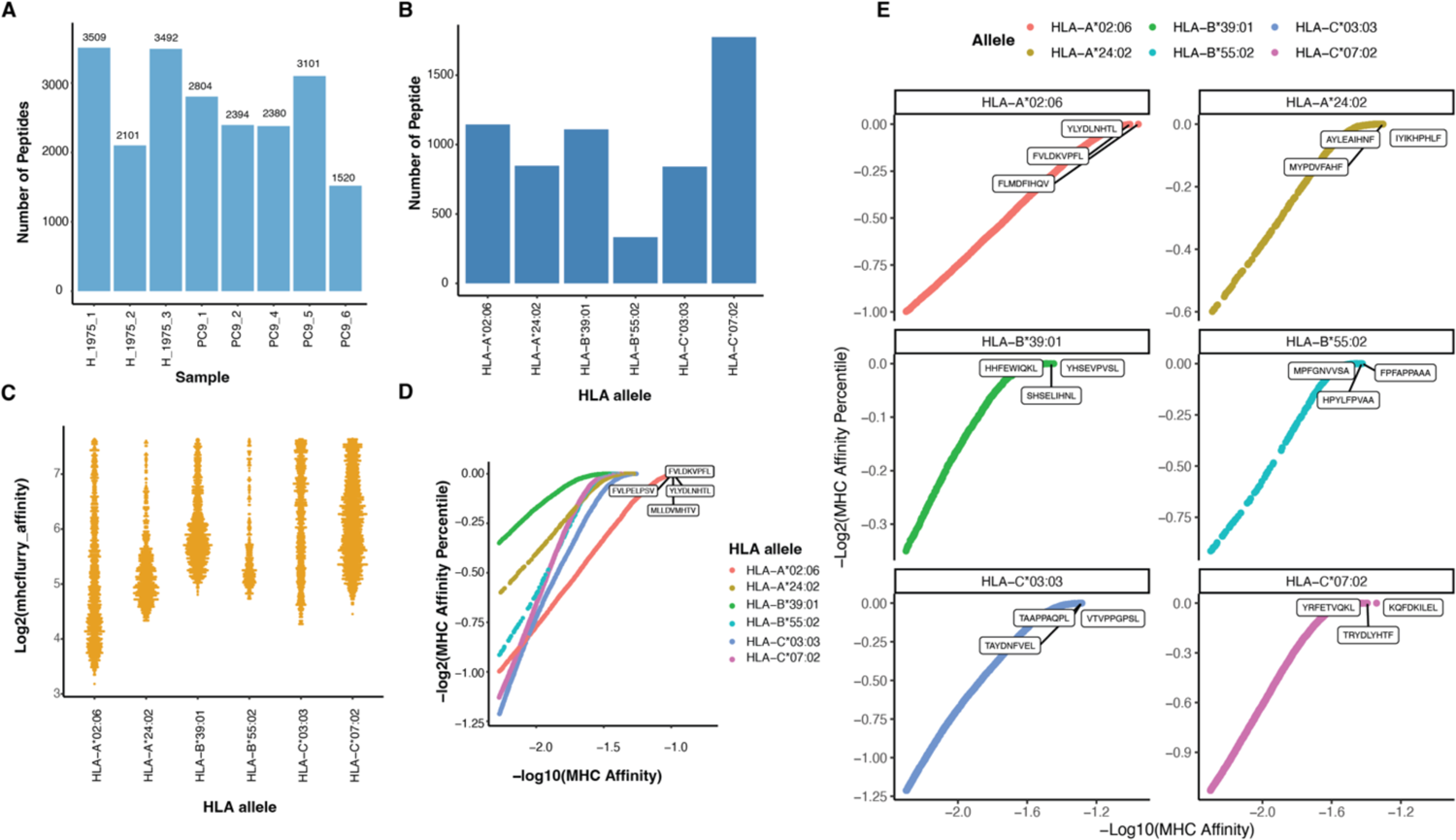
The characterization of peptides that bind specifically to particular MHC alleles. A. The count of peptides identified in class I immunopeptidome from various cancer cell lines and tumors. B. The count of peptides with HLA alleles has strong affinities. The cutoff of binding affinities C. The peptides’ binding affinities for each HLA allele with specific cutoff criteria for MHCflurry affinity (<200) and percentile(<2). D. Ranking Plot of Peptide-HLA Binding Affinities. The ranking plot visualizes the top 5 peptides exhibiting the strongest binding affinities. E. Peptide Binding Patterns by HLA Alleles. Top 5 peptides with robust binding affinities for each respective HLA allele shown.

### Case Study 4: Quantifying proteins in diverse brain tissue and related cells

To showcase the reliably and repeatedly identified proteins in common human specimen and cell line models in neurodegenerative diseases research using MS-proteomics, we compiled multiple proteomics datasets generated in-house sourced from a wide range of brain-related tissue and cellular contexts, encompassing human frontal cortex tissue, plasma, cerebrospinal fluid (CSF), skeletal muscle biopsies, as well as human iPSC-derived neuronal, microglial, and astrocytic cells. We assessed the protein count within each of these samples (Figure 6A). The samples derived from human tissues exhibited a comparatively lower number of detected proteins. Particularly, the plasma and CSF samples displayed the most modest protein counts, typically hovering around 3000 proteins. We performed UMAP on this dataset with the aim of examining data clustering patterns (Figure 6B). The iPSC microglial and astrocytic cells demonstrated a distinct clustering pattern, forming a cohesive cluster. In contrast, the iPSC neuron cells exhibited a clustering pattern more closely aligned with that of the muscle tissue samples.

**Figure 6.**
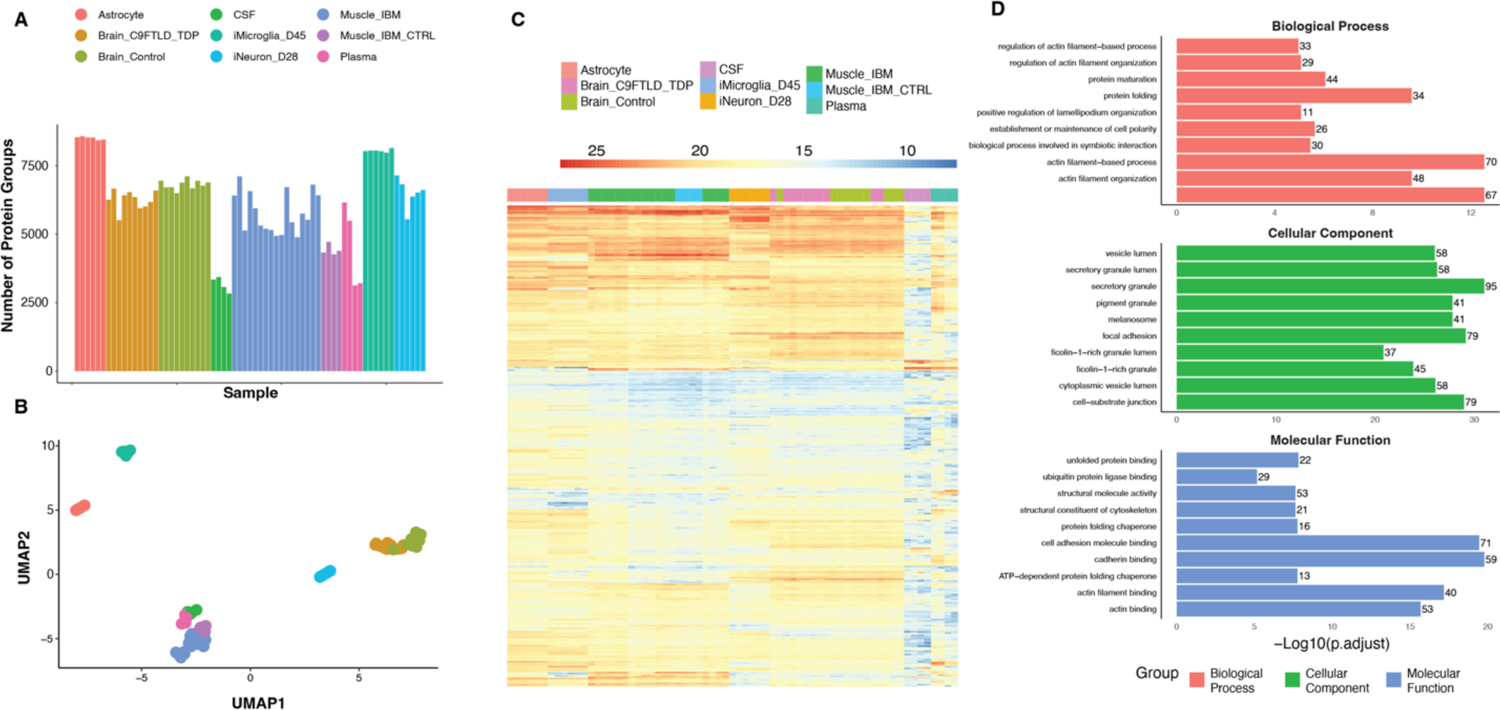
Analysis of proteomic datasets across diverse brain tissue and related cells A. The overview of the protein numbers of various tissue and cellular sources, including human brain samples, plasma, cerebrospinal fluid (CSF), muscle cells, and human induced pluripotent stem cell (iPSC)-derived neuronal, microglial, and astrocytic cells. B. The UMAP plot. C. Comprehensive heatmap of protein expression patterns across all detected proteins in the datasets D. Focused heatmap highlights a subset of 356 proteins consistently detectable across all samples.

Subsequently, our objective was to elucidate the protein expression patterns across the entirety of the samples. To achieve this, we generated a heatmap focusing exclusively on the subset of proteins that were consistently detectable across all samples, totaling 399 proteins (Figure 6C). Notably, our observations revealed a distinct subset of proteins exhibiting particularly pronounced expression within specific cellular contexts. Remarkably, this subset of approximately 300 proteins exhibited a discernible capacity to effectively distinguish between different cell types based on the clustering patterns observed in the heatmap. We also show the pathway analysis of these proteins that were consistently detectable across all samples (Figure 4D). The analysis of these datasets serves as a valuable resource for researchers seeking to identify benchmark proteins characteristic of distinct cell types and tissues.

## Discussion

MS has emerged as a pivotal technique in the realm of proteomics, offering a comprehensive approach for the identification and characterization of proteins. The versatility of MS extends its utility to diverse fields, including personalized medicine, systems biology, and various biomedical applications. By integrating MS with distinct proteomics methodologies such as AP-MS, immunopeptidomics, and total protein proteomics, researchers can delve into intricate aspects of biological systems. One of the remarkable capabilities of ProtPipe is its capacity to unravel multiple intricate modalities of proteomics, shedding light on the interconnectedness of cellular processes. AP-MS, in particular, has revolutionized the study of PPIs, enabling systematic exploration of complex biological networks. These insights are indispensable in understanding cellular processes, disease mechanisms, and facilitating drug discovery.

In the era of precision medicine, the advent of immunopeptidomics has been transformative. The comprehensive characterization and quantification of peptides presented by MHC on cell surfaces have unveiled crucial information for antigen presentation, immune recognition, and vaccine and adoptive immunotherapy development. The identification of tumor-specific antigens, viral epitopes, and the impact of diseases on the immunopeptidome pave the way for personalized immunotherapies and targeted vaccines against a spectrum of diseases. The sheer volume and intricacy of proteomics and peptidomics datasets pose significant challenges in data analysis. Accurate identification and quantification of proteins or peptides, coupled with the exploration of their biological functions in downstream analysis, demand meticulous data handling.

Imputation constitutes a pivotal technique within proteomics data analysis, employed to estimate and fill in missing values within datasets[26]. Missing values in DDA are commonly resulting from missing at random (MAR) due to the technical constraints [27]while those in DIA, Missing Not At Random (MNAR), are commonly due to the peptides’ low signal intensity, which is below the detection threshold[28].

Imputation methods are designed to rectify these gaps by approximating the absent values, thereby enhancing the comprehensiveness of datasets and diminishing the potential bias introduced by incomplete data. The imputation of missing values within the dataset extends the range of peptides available for analysis, resulting in an increased ‘yield’ for the experiment and, consequently, heightened sensitivity. Nevertheless, when these imputed values are not accurate, there is a potential for this method to yield incorrect experimental outcomes. For DIA data, we performed zero imputation with the assumption that missing values indicate non-detectable or absent protein abundance. Conversely, for Data-Dependent Acquisition (DDA) data, to mitigate the risk of false positive identifications, we recommend excluding samples with missing values.

Median normalization hinges on the assumption that most proteins undergo minimal alteration in their abundance levels when transitioning from one sample to another. Furthermore, any systemic bias that might be present is presumed to impact all proteins in a uniform manner. By employing this strategy, Median Normalization aims to align protein abundance values based on a central reference point, represented by the median. This approach, while effective in addressing systematic discrepancies, may not account for more nuanced variations specific to individual proteins.

The MHC molecules assume a pivotal role in introducing peptides to T cells, a fundamental step in immune recognition. This underpins their importance in orchestrating immune responses. In this context, various computational tools such as NetMHCpan[29], MHCflurry[30], and analogous resources come into play. These tools proficiently predict the binding affinity of peptides to MHC molecules, thereby aiding in the identification of neoantigen candidates poised for interaction with T cells. MHCflurry employs machine learning algorithms to train predictive models based on extensive experimental data. Through the adept utilization of intricate neural networks, MHCflurry effectively models the intricate interplay between peptides and MHC molecules, thereby facilitating precise and robust binding predictions. The tool’s prowess extends across various MHC alleles, enhancing its utility in diverse contexts and ensuring its accuracy across a spectrum of immunological scenarios.

In response to these complexities, we have developed ProtPipe, a multifunctional data analysis pipeline meticulously designed to streamline and automate the processing and analysis of high-throughput proteomics and peptidomics datasets. The pipeline encompasses a suite of features, including data quality control, sample filtering, and normalization. These aspects collectively contribute to the reliability and robustness of downstream analysis. ProtPipe offers researchers a spectrum of downstream analysis capabilities. It facilitates the identification of differential abundance proteins or peptides, enabling insights into the variation of protein expression across different conditions. Moreover, it empowers researchers with pathway enrichment analysis, enabling the exploration of functional implications of identified proteins. The protein-protein interaction analysis feature allows for the examination of intricate networks underlying biological systems. Additionally, it provides the valuable capability of MHC1-peptide binding affinity prediction. Importantly, ProtPipe is not only a robust tool but also an accessible one. It is an open-source software solution, readily available to the scientific community at <u>NIH-CARD/ProtPipe</u>. A web interface is under parallel development.

While ProtPipe offers a comprehensive suite of features to facilitate proteomics data analysis, it is essential to acknowledge its limitations. Firstly, the software’s performance may be influenced by the quality and completeness of the input data. Variations in data quality, such as missing values or outliers, can impact the accuracy of results, and while ProtPipe includes imputation and filtering strategies, addressing extreme data irregularities may require additional preprocessing steps. Secondly, ProtPipe’s versatility in accommodating multiple quantification tools is a strength, but it also introduces potential complexities in integrating and comparing results from different software packages. Researchers must exercise caution when interpreting data that combines outputs from various sources. Additionally, the current version focuses on certain aspects of proteomics data analysis, such as differential abundance and protein-protein interaction analysis. Future iterations may consider expanding functionalities to cover a broader spectrum of proteomics research needs. Lastly, while it strives for user-friendliness, its effectiveness is contingent on users’ familiarity with proteomics principles and data analysis concepts.

Thus, researchers with limited proteomics expertise may still face a learning curve when utilizing the software. Overall, acknowledging these limitations is crucial for researchers to make informed decisions when applying Protpipe to their specific proteomics investigations.

In conclusion, the marriage of MS with advanced proteomics approaches has propelled our understanding of protein function, cellular signaling, and complex biological systems. However, the magnitude and intricacy of the data generated necessitate sophisticated data analysis tools. ProtPipe, as presented here, represents a significant contribution to this field. Its multifunctional capabilities empower researchers to extract meaningful insights from high-throughput proteomics and peptidomics datasets efficiently and reliably, thus advancing our quest for a deeper understanding of intricate biological phenomena.

## Acknowledgement

This research was supported in part by the Intramural Research Program of the NIH, National Institute on Aging (NIA), National Institutes of Health, Department of Health and Human Services; project number ZIAAG000534, as well as the National Institute of Neurological Disorders and Stroke. M.P. is funded by NIH (RF1 NS120992, U54 NS123743). We thank the NIH HPC system (*Biowulf*, http://hpc.nih.gov) for making this work possible.

## Data availability

The MS raw files of case study 4 are deposited to the ProteomeXchange Consortium via the PRIDE [31]partner repository with the dataset identifier PXD047657. The MS raw files in case study 1, 2, and 3 are referred in original publications. Demo data can be found in the Protpipe GitHub (https://github.com/NIH-CARD/ProtPipe/tree/main).

## Author contributions

Z.L., and C.W. contributed equally to the general formulation and layout of the manuscript. A.B.S., M.A.N. and Y.A.Q. conceived and supervised the study. Z.Y., Y.A.Q., and M.A.N. designed and conceptualized this pipeline. Y.H., J.R., C.B., and S.K. carried out the mass spectrometry experiments. P.F., L.P., M.P. and D.W.D., supervised and collected the human muscle and/or brain samples. B.O. and N.S. collected the human CSF samples. M.R.C., and M.E.W. supervised and carried out the iPSC-derived neurons differentiation and collection. Z.L., and C.W. performed proteomics data analysis and integrative bioinformatics analyses. C.W., and N.J. developed the Singularity image. S.S. developed the web app. Z.L prepared the manuscript. All authors contributed to writing of the manuscript, as well as reviewed and accepted the manuscript.

---

**Supplementary figure 1.**
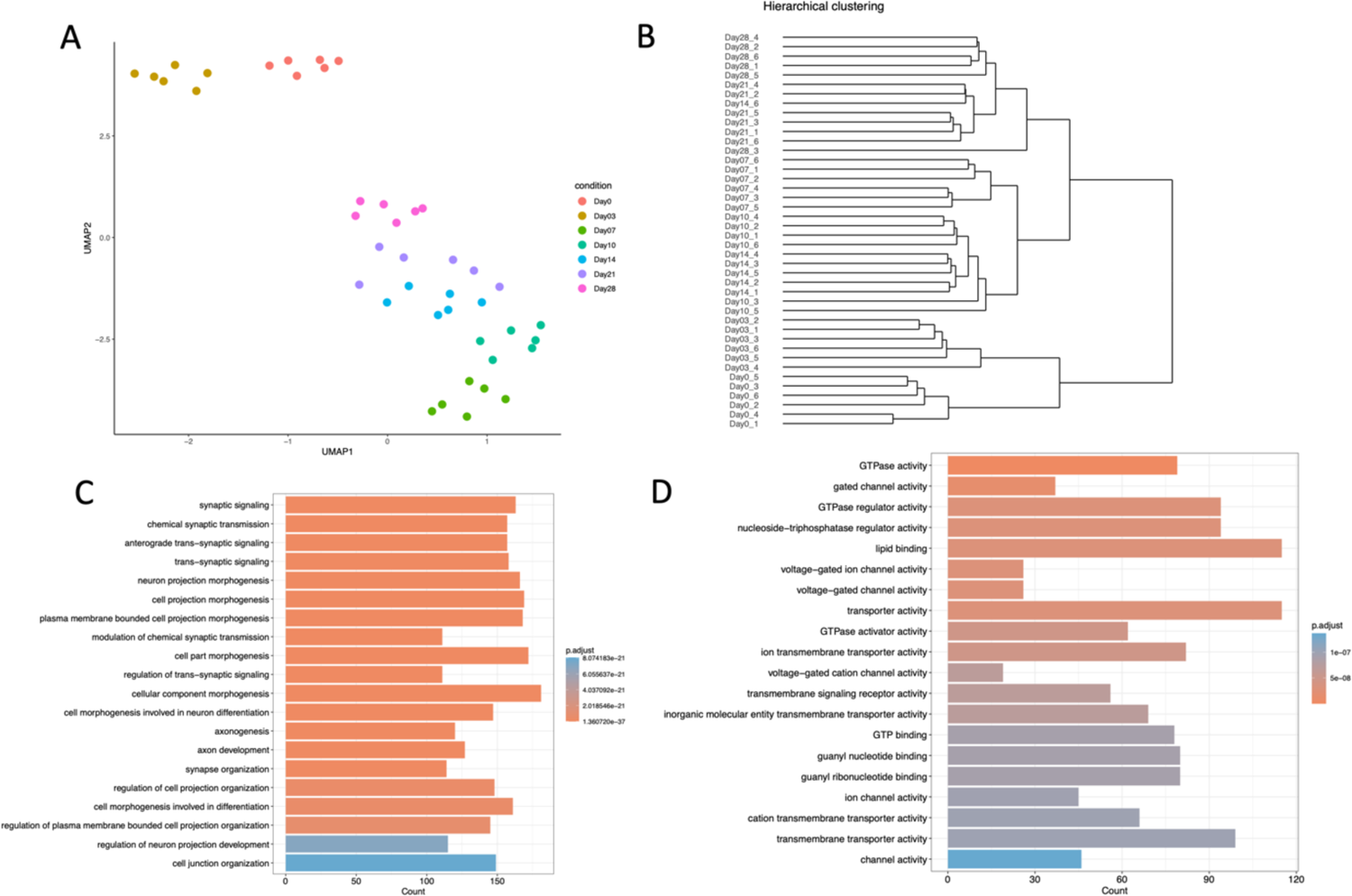

**Supplementary figure 2.**
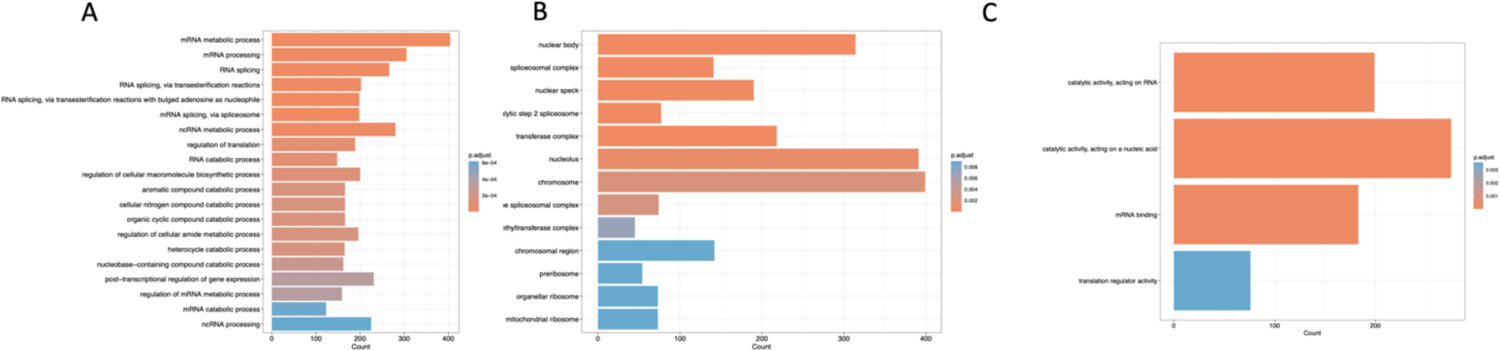

**Supplementary figure 3.**
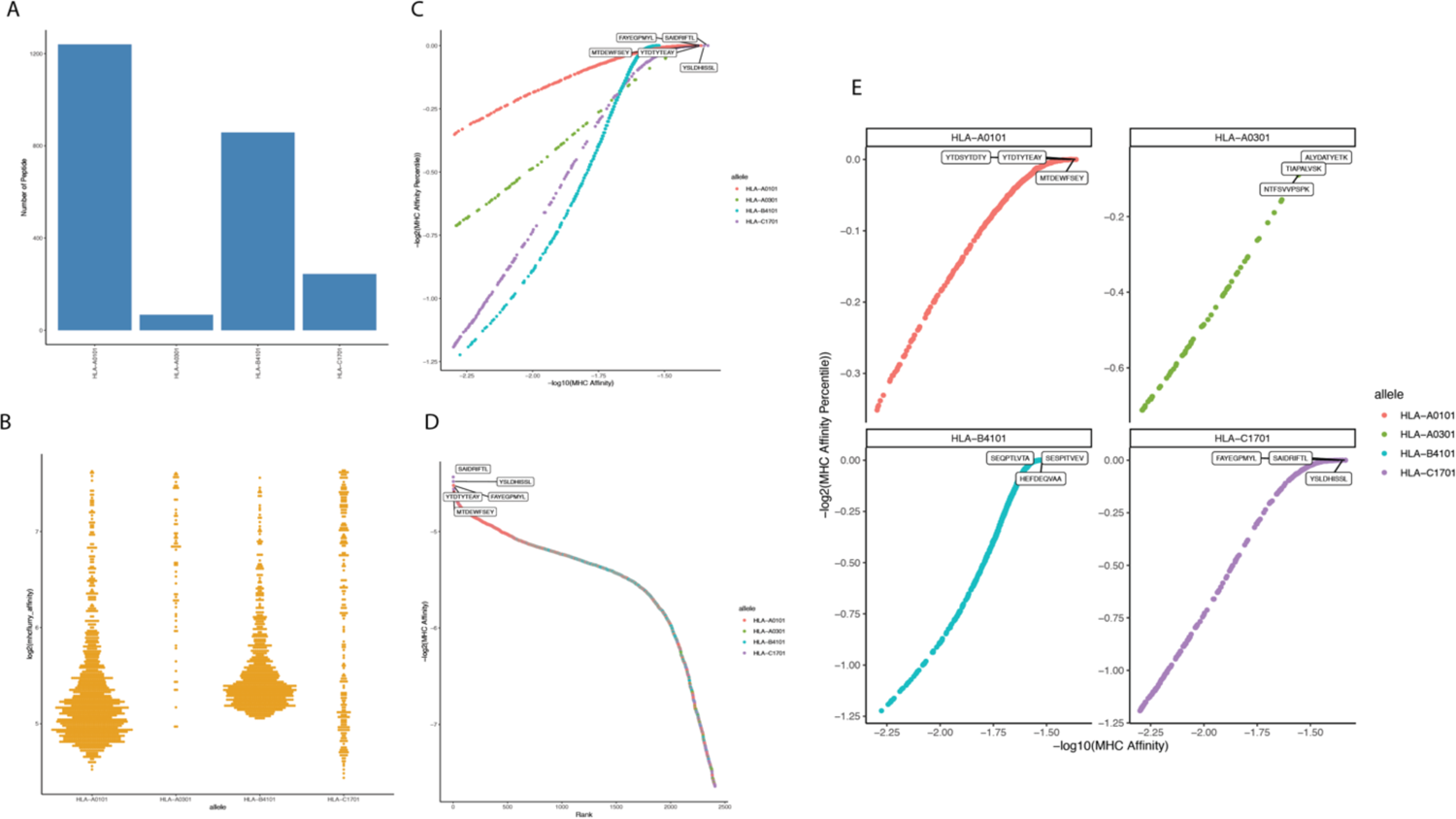

